# Response-driven Serial Dependence: When Response Mode Consistency Matters More than Shared Memory Encoding

**DOI:** 10.1101/2025.08.26.672319

**Authors:** Jiao Wu, Halid Oğuz Serçe, Zhuanghua Shi

**Affiliations:** Department Psychologie, Ludwig-Maximilians-Universität München, Munich, Germany; Bahçeşehir University, Istanbul, Turkey

**Keywords:** serial dependence, sequential biases, motor response, decisional carryover

## Abstract

Serial dependence - the bias from recent experience on present response - is often linked to shared memory representations. Yet, it remains unclear whether this bias across tasks when stimulus features are shared but response modes differ. To test this, we interleaved temporal reproduction and bisection tasks using a post-cue design that held duration encoding constant while varying motor output across trials. Using structural equation modeling (SEM), we dissociated perceptual (stimulus-driven) and decisional (response-driven) components of serial dependence. With consecutive same tasks, we replicated repulsive perceptual serial dependence and attractive decisional carryover. Critically, these effects vanished across tasks, despite identical stimulus processing – suggesting that consistent motor responses, not shared memory alone, drive sequential biases. Moreover, SEM uncovered repulsive perceptual influences that standard regression missed, highlighting its power to isolate overlapping effects. Together, these findings reveal that response-specific reactivation underpins serial dependence, pointing to motor-context binding as a key factor in temporal decision-making.

## Introduction

Imagine trying to judge whether a line you see is vertical or slightly tilted after viewing a series of tilted lines. Your judgment tends to lean toward the orientation you have most recently observed, reflecting a systematic bias caused by prior experience. This bias, known as *serial dependence* or sequential effect (Fischer & Whitney, 2014; Manassi et al., 2023; Pascucci et al., 2023), indicates how recent sensory history shapes current perceptual decisions. Serial dependence has been documented across various stimulus properties, including orientation, motion direction, position, color, shape, and numerosity (Cicchini et al., 2024; Pascucci et al., 2023). Crucially, such effects can occur at multiple cognitive levels - from early perceptual mechanisms (Burr & Cicchini, 2014; Fischer & Whitney, 2014; Kiyonaga et al., 2017; Liberman et al., 2016) to high-level functions involving memory and decision-making (Fritsche et al., 2020; Hahn & Wei, 2024; Hajonides et al., 2023; Pascucci et al., 2019, 2023; Sheehan & Serences, 2022; Zhang & Lewis-Peacock, 2024).

Not just the perception of stimulus features, time perception is also subject to serial dependence (Cheng, Chen, Glasauer, et al., 2024; Glasauer & Shi, 2022; Li et al., 2023; Wang et al., 2023; Wiener et al., 2014). Unlike orientation or color, perceived duration lacks a dedicated sensory system, relying instead on distributed neural and cognitive processes (Ivry & Schlerf, 2008; Merchant et al., 2013). Interestingly, sequential biases in temporal judgments have been observed across various timing tasks, including duration reproduction and temporal bisection, even when the classical central tendency effect was carefully controlled for (Cheng, Chen, & Shi, 2024; Glasauer & Shi, 2022; Li et al., 2023). Intriguingly, though, magnitudes of sequential effects depend on task context or switching between different modalities or tasks (Bae & Luck, 2020; Cheng, Chen, & Shi, 2024; Cheng, Chen, Yang, et al., 2024; Li et al., 2023). For instance, the sequential effect was reduced or diminished when the task was switched across trials, requiring participants to reproduce either direction or duration based on a trial-by-trial cue (Cheng, Chen, & Shi, 2024).

These modality- and task-specific sequential effects suggest that post-perceptual processes, particularly working memory, play a key role in serial dependence (Bae & Luck, 2020; Bliss et al., 2017; Cheng, Chen, Yang, et al., 2024). Maintaining the same task-relevant information across trials (e.g., duration) can facilitate the integration of past and present stimuli in working memory and/or reactivation of subthreshold traces of recent memories (Bae & Luck, 2019; Barbosa & Compte, 2020), resulting in an attraction bias (Cheng, Chen, & Shi, 2024; Cheng, Chen, Glasauer, et al., 2024; Cheng, Chen, Yang, et al., 2024). Despite extensive research on serial dependence, most findings related to task-specific or modality-specific effects come from studies interleaving tasks with distinct features or modalities, such as direction versus time judgments (Cheng, Chen, & Shi, 2024; Cheng, Chen, Yang, et al., 2024), numerosity versus time judgments (Fornaciai et al., 2023; Togoli et al., 2021), or color versus direction tasks (Bae & Luck, 2020). Such distinct task pairings typically require participants to concurrently encode two separate stimulus attributes (e.g., motion direction and duration) into working memory. Selecting one task typically enhances the representation of its relevant feature while suppressing the representation of its irrelevant features, resulting in minimal or no serial dependence across tasks (e.g., Bae & Luck, 2020).

However, previous studies leave open the question of whether task-specific serial dependence emerges when interleaved tasks share identical features (e.g., duration) but differ only in their motor response requirements. This distinction is crucial for clarifying the cognitive bias of serial dependence because it isolates contributions of the response mode from feature-based memory representation. Prior research has shown that both the previous stimulus and response influence serial dependence, but separating their effects remains challenging (Moon & Kwon, 2022; Sadil et al., 2024). If serial dependence primarily relies on shared memory representations, interleaving two tasks involving the identical features should yield similar sequential biases despite differences in response mode. Conversely, if serial dependence depends on task-specific response reactivation, identical stimulus content might still yield differential sequential effects when response modes switch. Supporting the relevance of response modes, prior research, although not directly addressing serial dependence, has shown that estimation biases can differ with specific motor outputs (e.g., manual vs. vocal responses) even with the same task (Roach et al., 2017).

In the present study, we directly investigate this issue by interleaving two temporal tasks – duration reproduction and temporal bisection – that use identical duration stimuli but differ in their motor responses. Participants performed trials involving either reproducing a given interval by manual pressing (reproduction task) or categorizing intervals as short or long by binary responses (bisection task). A post-cue design was employed, where participants encoded and maintained the duration first and executed the required tasks only after a cue was presented, ensuring identical perceptual encoding across trials. This design isolates potential contributions of motor response from those related to perceptual and memory encoding. By examining serial dependence both within and across these tasks, we aimed to clarify the role of motor responses in shaping sequential effects in time perception.

## Methods

### Participants

We recruited 24 volunteers (16 females, 28.5 ± 4.49 years) for the experiment. The sample size was based on effect sizes, ranging from 0.596 to 1.84, reported in previous studies on serial dependence in duration reproduction tasks (Cheng, Chen, Yang, et al., 2024; Glasauer & Shi, 2022) and the effect size of 0.684 in a duration bisection task (Cheng, Chen, Yang, et al., 2024). To ensure sufficient power (80%) at an alpha level of 0.05, we used the smallest effect size (0.596), which yielded at least 19 participants according to G*Power calculation (Erdfelder et al., 1996). To be conservative, we recruited 24 participants to assess both task-relevant and potential task-irrelevant serial dependence. All reported normal or corrected normal vision, gave informed consent, and received monetary or course credit compensation for their participation. The Ethics Committee of the Department of Psychology of LMU approved the study.

### Apparatus and stimuli

The experiment took place in a soundproof, dimly lit room (2.0 cd/m^2^) with a table lamp placed behind a 19-inch CRT monitor (85 Hz refresh rate). The experiment was implemented using Matlab (Mathworks Inc.) and Psychtoolbox (Kleiner et al., 2007).

Duration stimuli were presented as Gabor patches displayed for a randomly selected duration from a uniform distribution of 500, 610, 740, 910, 1100, 1340, 1640, and 2000 ms, with equal spacing by 0.197 log units in the log scale. The geometric mean of the sampled durations was 1000 ms. The Gabor patches subtended 5°×5° in visual angle and had a spatial frequency of 0.5 cycles per degree. Their orientation was randomly chosen from -81° to 81°, in 18° increments. The sinusoidal gratings of Gabor had a peak contrast of 100% Michelson and were enveloped by a Gaussian hull with a standard deviation of 2.29° radius. All stimuli and task cues appeared against a gray background (13.18 cd/m^2^).

### Design and procedure

The experiment involved two tasks: duration *reproduction* and duration *bisection*. In the reproduction task, participants reproduced the duration of a presented stimulus as accurately as possible. In the bisection judgment task (Wearden, 1997), participants judged if the presented stimulus duration was shorter or longer than a standard duration (1 second), shown five times at the beginning of each experimental block.

Participants received task instructions only after the stimulus presentation following a post-cue design: a dot indicated the reproduction task, while left- and right-pointing arrows signaled the bisection task (see Figure 1). This design ensured consistent duration encoding across trials, independent of tasks, thus allowing us to assess if sequential effects generalized across task types.

**Figure 1.**
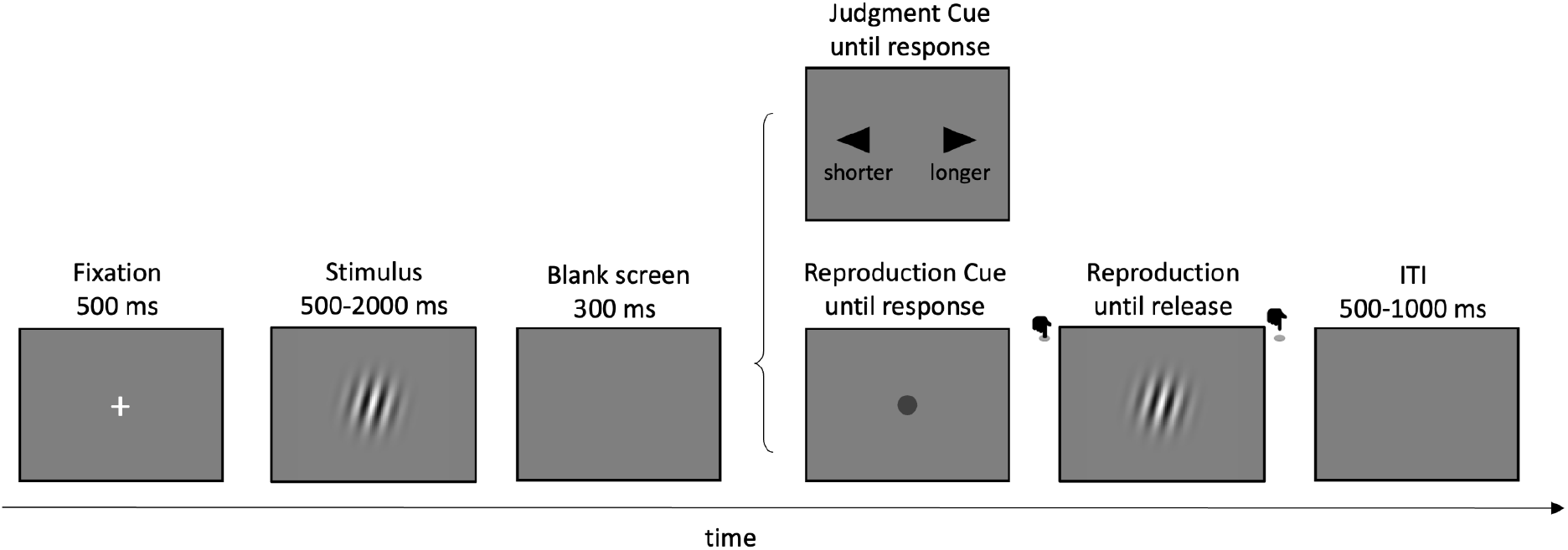
Schematic illustration of trial sequence. Each trial started with a 500-ms central fixation, followed by a Gabor patch displayed for a given duration (500, 610, 740, 910, 1100, 1340, 1640, or 2000 ms). After a 300-ms blank screen, a task cue appeared, informing the task for the current trial. In reproduction trials, observers pressed a key to reappear the same Gabor patch for the perceived duration. In bisection trials, they judged if the current duration was shorter or longer than the standard duration, shown multiple times at the start of each block. After the response, a random blank interval (500-1000 ms) was presented before the next trial.

The experiment comprised 20 blocks, each containing 32 trials, for a total of 640 trials. Each duration appeared 40 times per task type. Within each block, we randomized and balanced duration levels and task types, repeating each condition twice. To ensure equal representation, inter-trial transitions from trial *n*-1 to trial *n* equally covered all possible task combinations: reproduction followed by reproduction (RR), bisection judgment followed by reproduction (JR), reproduction followed by bisection judgment (RJ), and judgment followed by judgment (JJ).

Each trial started with a 500-ms central fixation, followed by a Gabor patch for a randomly selected duration. After a 300-ms blank screen, a task cue, either a dot (reproduction task) or left- and right-arrows (bisection task), appeared to indicate the task for the current trial. In the duration reproduction task, a dot cue remained visible until observers pressed a key. Upon keypress, the same Gabor patch reappeared, remaining on the screen until the key was released. The keypress duration indicated the reproduced duration. In the bisection judgment task, two directional arrows appeared, prompting observers to indicate via keypress whether the perceived duration was ‘shorter’ or ‘longer’ than the 1-second standard. After each response, a blank inter-trial interval (ITI) of 500-1000 ms preceded the next trial.

Prior to the formal experiment, observers practiced a block of 32 trials to familiarize themselves with the tasks. During practice, they received accuracy feedback lasting for 500 ms. For reproduction practice trials, feedback displayed ‘Too short’, ‘Well done’, or ‘Too long’ when the relative reproduction error ([Reproduced duration - Actual duration]/Actual duration) fell below -30%, within [-30%, 30%], or above 30%, respectively. In bisection trials, feedback indicated ‘Correct’ or ‘Incorrect’ based on whether the response matched the comparison between the presented duration and the 1-second standard.

### Statistical Analysis

Considering that reproduced durations are influenced by both *central tendency* and *serial dependence* (approximately linear; Glasauer & Shi, 2022), we treated the reproduction bias (i.e., reproduced minus physical duration) as linearly driven by both the current and prior stimuli. Importantly, serial dependence combines two distinct processes: (a) sensory carryover, a bias from the prior stimulus; and (b) decisional carryover, an attractive bias toward the prior response. However, the direction of sensory carryover is rather mixed. Some found repulsive biases (Fritsche et al., 2017; Pascucci et al., 2019; Sadil et al., 2024), while others showed attractive biases (Cheng, Chen, & Shi, 2024; Cheng, Chen, Yang, et al., 2024; Fischer & Whitney, 2014; Glasauer & Shi, 2022). These mixed findings could be a methodological artifact: when both prior stimulus and prior response are considered, while sharing covariance due to their inherent correlation, potentially leading to multicollinearity. Consequently, the true stimulus-based sensory carryover of serial dependence may be masked or canceled out by decisional carryover of prior responses with standard regressions.

To this end, we applied structural equation modeling (SEM), which offers several key advantages over standard regression models. It allows explicit modeling of latent structures, enabling causal modeling via directed paths (see Figure 2): current stimulus → current response; prior stimulus → bias (direct sensory carryover), and prior report → current response (decisional carryover). This model also includes an indirect sensory carryover: prior stimulus → prior report → current response, mirroring theoretical processes of sensory versus decisional serial dependencies. Unlike conventional regression, SEM incorporates measurement error, enhancing parameter reliability. By explicitly modeling the covariance between the prior stimulus and the prior response, SEM alleviates multicollinearity issues. To our knowledge, no prior study has used SEM to dissect sensory versus decisional carryover in this manner.

**Figure 2.**
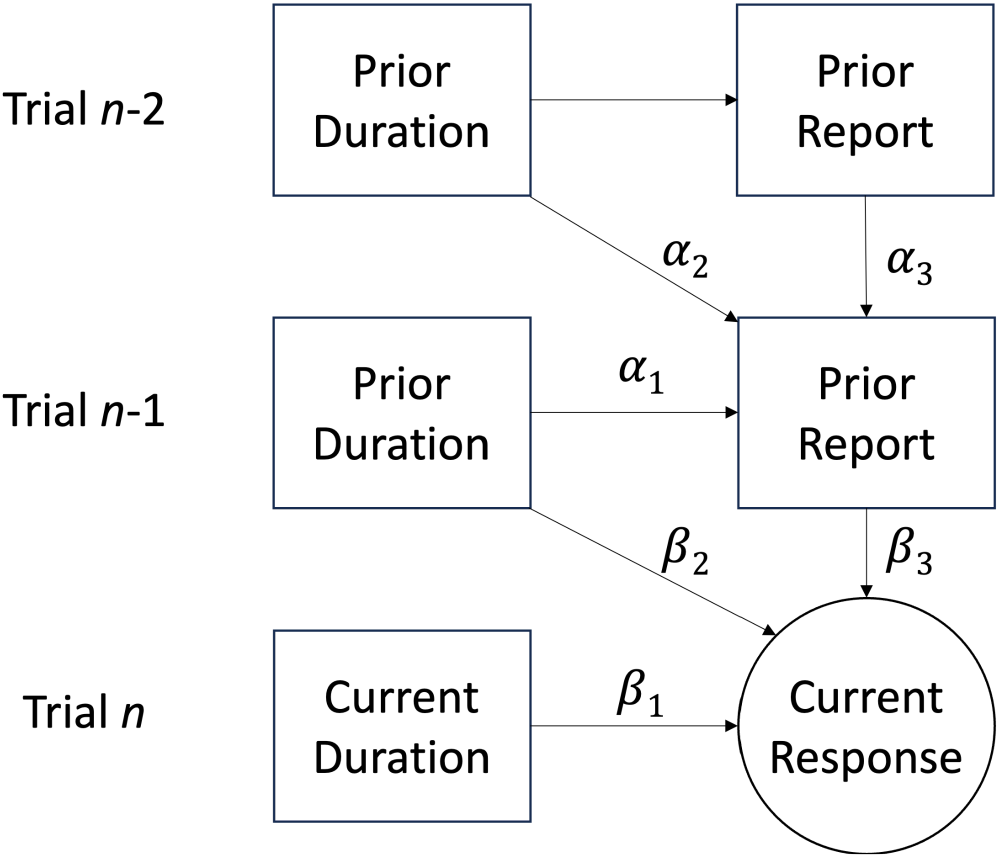
The schematic of structural equation modeling (SEM). The model specifies directed paths: current stimulus → current response (β_1_); prior stimulus → bias (direct sensory carryover, β_2_), and prior report → current response (decisional carryover, β_3_). This model also includes an indirect sensory carryover: prior stimulus → prior report → current response (α_1_*β_3_), mirroring theoretical processes of sensory versus decisional serial dependencies. The causal influences can be traced to earlier trials (e.g., trials n-2).

Specifically, we constructed structural paths to parse the causal influences of current duration (β_1_), prior duration (*β*_2_), and prior report (short/long category) in trial n-1 (*β*_3_) on the current response (Figure 2). Such causal influences can be traced back to earlier trials. To unify the two task types, we assumed that decisional carryover mainly stems from the coarse categorization of the prior report (short or long). In the bisection task, this categorization was explicit; in the reproduction task, we treated it as implicit, based on how responses compared to their mean reproduction. To account for the scalar property in duration estimation (Gibbon et al., 1984; Ren et al., 2021), we modeled time duration on a log-scale.

We applied SEM to both tasks using the *lavaan* package in R for model specification, estimation, and evaluation (Rosseel, 2012). The categorical variables (e.g., the prior report and the bisection response) were treated as ordinal endogenous variables. *Lavaan* automatically applies a threshold SEM using the diagonally weighted least squares with mean-variance adjustment (WLSMV) estimator. For binary responses in the bisection task, this is formally equivalent to a probit regression. That is, *lavaan* models an underlying continuous latent variable Y^*^ (the latent response) with thresholds, and the probability of a positive (“long”) response is *P*(*Y*^*∗*^ > 0|*X*) = *Φ*(*X*^*T*^*β*), where *Φ* is the standard normal cumulative distribution function. This model is analogous to the conventional measure estimating sequential effect in binary judgment tasks — the probabilistic choice model (Feigin, Baror, et al., 2021; Feigin, Shalom-Sperber, et al., 2021; Li et al., 2023), which takes into account current stimuli, prior stimuli, and prior report together (but potentially encountering multicollinearity issues). To compare with the conventional probabilistic choice model, we also conducted the conventional approach (see Appendix).

## Results

The missed trials in the bisection task and trials with extreme reproduced durations (i.e., shorter than 50 ms or exceeding 5 secs) were excluded from the analysis (0.40% in total). As for the four task combinations for trial *n*-1 and *n* (i.e., RR, JR, RJ, and JJ), the inter-trial transitional probability was equal but with a slight cross-subject variance due to randomization (25.00% ± 1.17%).

### Reproduction task

We applied SEMs to individual participants on each prior task type (reproduction vs. bisection). The model provided excellent fits to the data, with mean χ^2^(3) values of 3.07 for the RR condition and 3.20 for the JR condition (*p*s > 0.508). The mean Goodness of Fit Indices (GFIs) were 0.988 and 0.989 for the RR and JR, respectively. These indices meet conventional thresholds, supporting acceptable model performance. Table 1 presents the mean coefficients from the SEM models, along with related statistical results (see also Figure 3).

**Table 1.** Estimates of the structural equation modeling (SEM) for the reproduction task.

| | RR<br>($M \pm SE$) | JR<br>($M \pm SE$) | $t$ | $df$ | $p$ | Cohen's $d$ | $BF_{10}$ |
| --- | --- | --- | --- | --- | --- | --- | --- |
| $\beta_1$ | -0.421 ± 0.050 *** | -0.409 ± 0.050 *** | -0.751 | 23 | .460 | -0.153 | 0.277 |
| $\beta_2$ | -0.212 ± 0.037 *** | -0.024 ± 0.041 | -4.357 | 23 | < .001 | -0.889 | > 100 |
| $\beta_3$ | 0.097 ± 0.018 *** | 0.010 ± 0.012 | 5.096 | 23 | < .001 | 1.040 | > 100 |
| $\beta_4$ | 0.251 ± 0.034 *** | 0.025 ± 0.039 | 5.181 | 23 | < .001 | 1.058 | > 100 |
| $\beta_5$ | 0.038 ± 0.013 ** | 0.001 ± 0.013 | 3.047 | 23 | .006 | 0.622 | 7.765 |

Note: The parameter $\beta_4$ represents the indirect mediating effect ( $\beta_4 = \alpha_1 * \beta_3$ ) and $\beta_5$ represents the net serial dependence (net effect from prior duration: $\beta_5 = \beta_2 + \alpha_1 * \beta_3$ ). The statistics, including $t$ , $p$ , $df$ , Cohen's $d$ , and Bayes Factor, are based on the comparison between the relevance conditions of prior task type (i.e., RR vs JR). Stars \*\* and \*\*\* represent $p < .01$ and .001, respectively, for statistical significance compared to zero.

**Figure 3.**
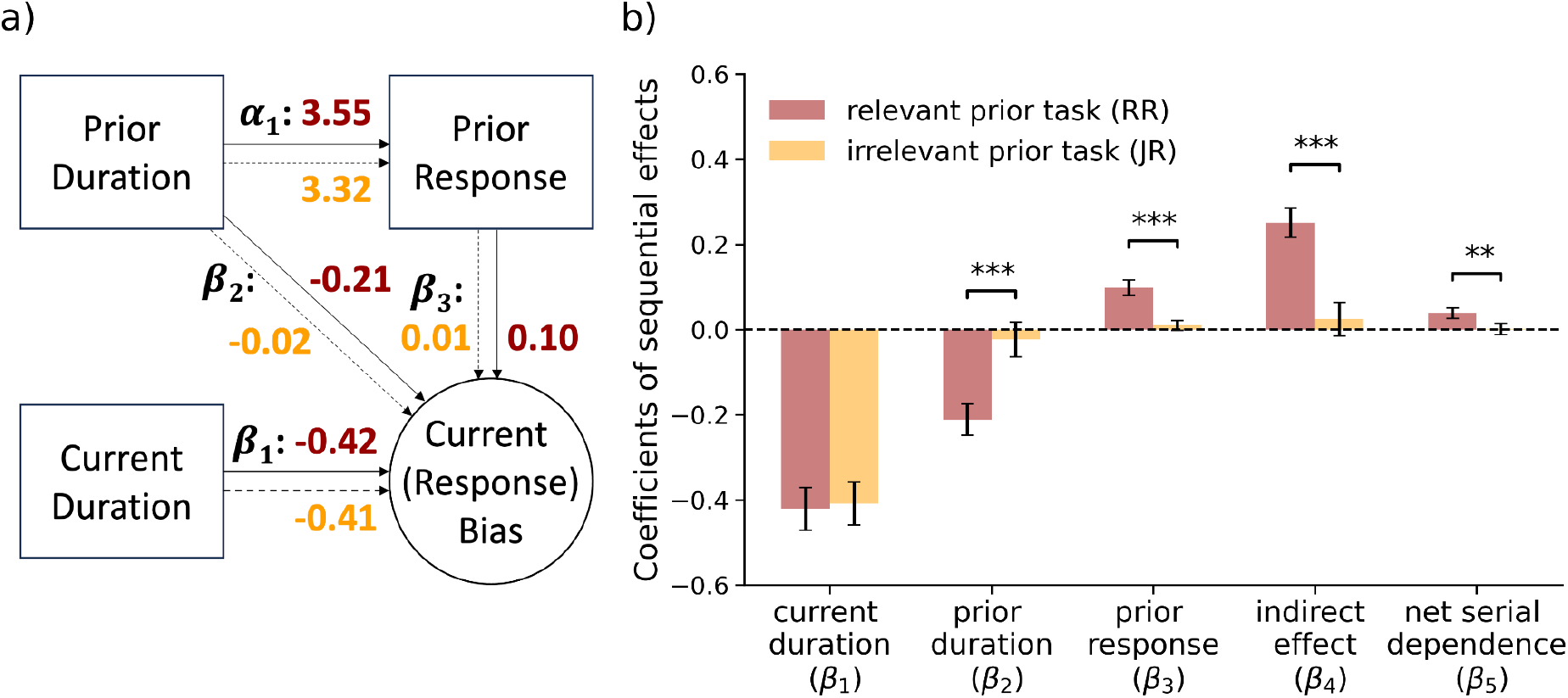
Reproduction task performance. (**a)** Estimated coefficients in SEM for the reproduction task. The first row shows the predictors from a prior trial, and the second shows predictors in the current trial. The solid lines and coefficients in dark red show the parameter estimates, which describes the strength and direction for the relationships between variables, for the task-relevant (RR) trials. The dashed lines and coefficients in orange depict the estimates for the task-irrelevant (JR) trials. (**b)** Parameter estimates for prior task types, predicted by the structural equation modeling (SEM). The error bar shows standard errors across participants. *** p <.001, ** p <.01.

#### Direct Effects

##### The central tendency effect (*β*_1_)

The current log-transformed duration showed a significant negative linear relationship with reproduction bias in both RR (-0.421) and JR (-0.409) conditions (Table 1), confirming the classic central tendency biases. However, the central tendency biases were comparable between the two conditions (*p* =.460).

##### Perceptual serial dependence (*β*_2_)

While the prior duration had a significantly negative slope (-0.212) on the current bias in the RR task-congruent sequence (*p* <.001), it did not show significance in the JR task-incongruent sequence (*p* =.567). The negative slope shows that longer prior durations caused a negative bias, reflecting a common repulsion effect. The difference between the two conditions was also significant (*p* <.001, Figure 3b), suggesting that prior task-relevance is an important factor in perceptual serial dependence.

##### Decisional carryover (*β*_3_)

Similar to perceptual serial dependence, decisional carryover also exhibited differential effects in two conditions. But instead of negative repulsion, the carryover was a significantly positive attraction (0.097) in the RR condition (*p* <.001), but non-significant in the JR condition (*p* =.408). The difference between the two conditions was also significant (*p* <.001). The pattern suggests that decisional carryover occurs only with relevant tasks.

#### The Indirect Effect

One advantage of using SEM is that it allows us to estimate the indirect effect. A significant indirect effect emerged via the pathway from prior duration → prior report → reproduction bias (*β*_*4*_ *= α*_*1*_ *\* β*_3_) in the RR condition (*p* <.001) but failed to show any effect in the JR condition (*p* =.540). The difference between the two conditions was also significant (*p* <.001).

#### The Net Serial Dependence

The net effect from prior duration (*β*_*5*_ *= β*_*2*_ *+ α*_*1*_ *\** β_3_) was consistent with the serial dependence measured by conventional regression (see Appendix). The net serial dependence showed significance in the RR (*p* =.006) but not the JR (*p* =.943) condition. The difference between the task-relevance conditions was also significant (*p* =.006). This further suggests that conventional serial dependence could be the cancellation of perceptual serial dependence and decisional carryover.

### Bisection Judgment task

The generalized SEMs (probit models) for the bisection task also provided excellent fits to the data, with mean χ^2^(3) values of 2.62 for the RJ condition and 1.93 for the JJ condition (*p*s > 0.542). The mean Goodness of Fit Indices (GFIs) were 0.960 and 0.975 for the RJ and JJ conditions, respectively. Table 2 presents the mean coefficients from the generalized SEM models, along with related statistical results (see also Figure 4).

**Table 2.**
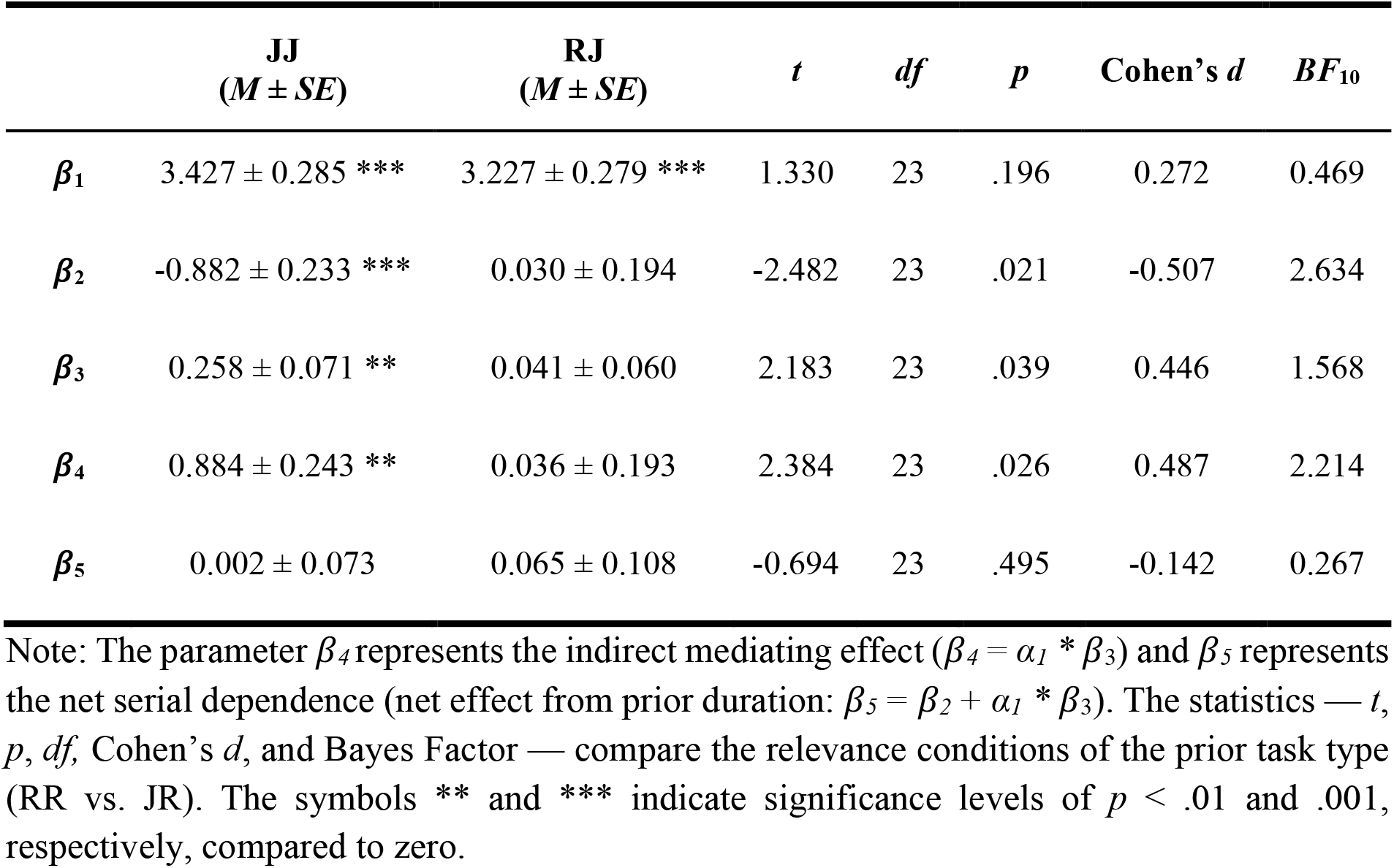
Estimates of the structural equation modeling (SEM) for the judgment task.

| | JJ<br>($M \pm SE$) | RJ<br>($M \pm SE$) | $t$ | $df$ | $p$ | Cohen's $d$ | $BF_{10}$ |
| --- | --- | --- | --- | --- | --- | --- | --- |
| $\beta_1$ | $3.427 \pm 0.285$ *** | $3.227 \pm 0.279$ *** | 1.330 | 23 | .196 | 0.272 | 0.469 |
| $\beta_2$ | $-0.882 \pm 0.233$ *** | $0.030 \pm 0.194$ | -2.482 | 23 | .021 | -0.507 | 2.634 |
| $\beta_3$ | $0.258 \pm 0.071$ ** | $0.041 \pm 0.060$ | 2.183 | 23 | .039 | 0.446 | 1.568 |
| $\beta_4$ | $0.884 \pm 0.243$ ** | $0.036 \pm 0.193$ | 2.384 | 23 | .026 | 0.487 | 2.214 |
| $\beta_5$ | $0.002 \pm 0.073$ | $0.065 \pm 0.108$ | -0.694 | 23 | .495 | -0.142 | 0.267 |

Note: The parameter $\beta_4$ represents the indirect mediating effect ( $\beta_4 = \alpha_I * \beta_3$ ) and $\beta_5$ represents the net serial dependence (net effect from prior duration: $\beta_5 = \beta_2 + \alpha_I * \beta_3$ ). The statistics — $t$ , $p$ , $df$ , Cohen's $d$ , and Bayes Factor — compare the relevance conditions of the prior task type (RR vs. JR). The symbols \*\* and \*\*\* indicate significance levels of $p < .01$ and $.001$ , respectively, compared to zero.

**Figure 4.**
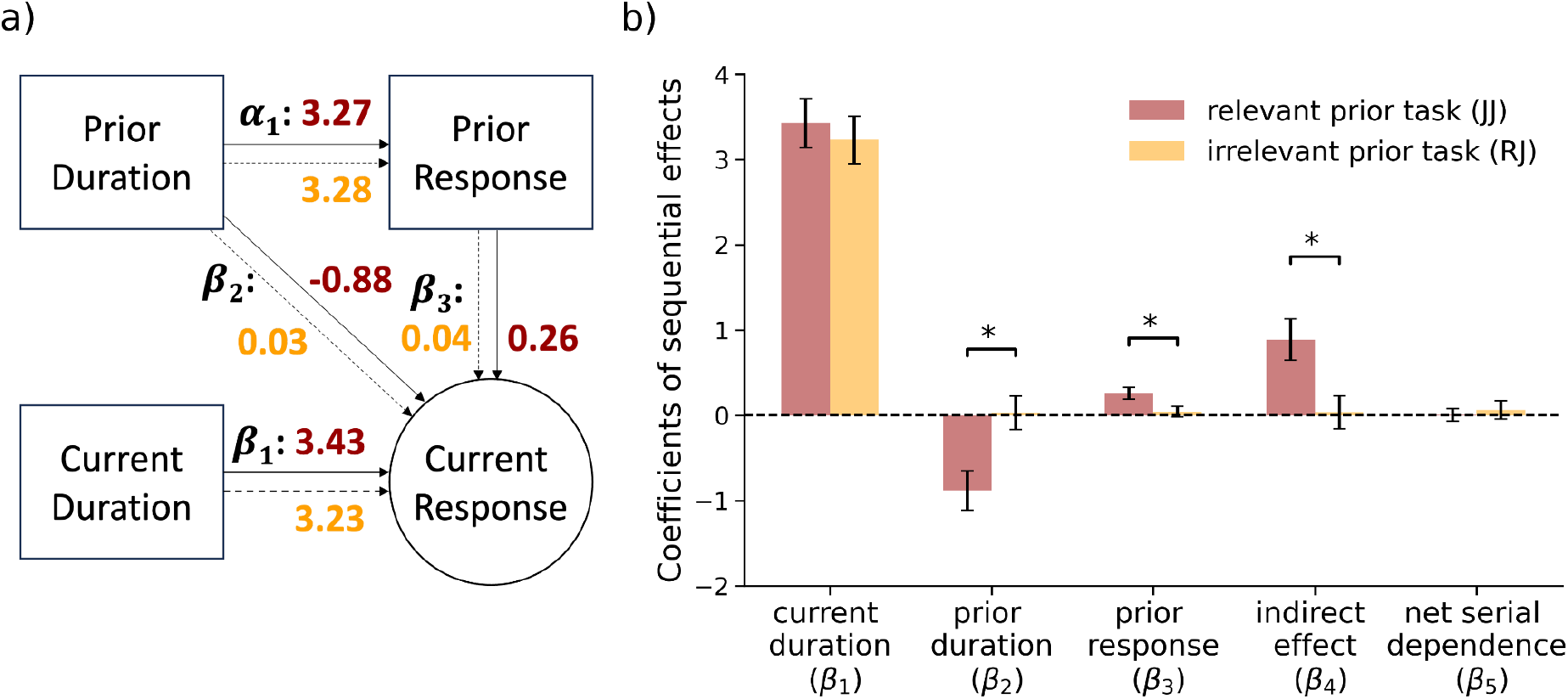
Judgment task performance. (**a)** Estimated coefficients in SEM for the judgment task. The first row shows the factors from a prior trial, and the second shows factors in the current trial. The task-relevant coefficients (JJ) are with solid lines and in dark red, while the task-irrelevant coefficients (RJ) are with dashed lines and in orange. (**b)** Parameter estimates for prior task types predicted by the structural equation modeling (SEM). The error bar shows the standard error across participants. * p <.05.

It is important to note that the coefficients *β*_1_, *β*_2_, and *β*_3_ correspond conceptually to central tendency, perceptual serial dependence, and decisional carryover, respectively, they do *not* directly measure those effects in the observed data scale. Rather, they quantify the impact of predictors on the *latent* response variable Y^*^ (a z-score scale), not on the raw behavioral bias. Specifically, *β*_1_ expresses how a one-unit increase in log-transformed current duration shifts the latent decision propensity Y^*^ toward “long” by *β*_1_ standard deviations. Table 2 shows that both RJ and JJ had comparable *β*_1_ (*p* =.196).

The coefficients *β*_2_ and *β*_3_ similarly reflect how prior stimulus and prior response influence that latent decision tendency. Similar to the reproduction task, the prior duration had a significantly negative *repulsion* on the latent response variable Y^*^ in the JJ task-congruent sequence (*p* <.001); it did not show significance in the RJ task-incongruent sequence (*p* =.882). The difference between the two conditions was also significant (*p* =.021, Figure 4b). The decisional carryover on the latent variable also exhibited differential effects in two conditions. But instead of negative repulsion, the carryover was significantly positive (i.e., attractive) in the JJ condition (*p* =.001), but not in the JR condition (*p* =.506). The difference between the two conditions was also significant (*p* =.039). The pattern suggests that decisional carryover in the latent space occurs only with relevant tasks.

The coefficient *β*_4_, reflected the indirect effect via the pathway from prior duration → prior report → reproduction bias, showed significance in the JJ (*p* =.001) but not RJ (*p* =.855) sequence. The comparison between task-congruence conditions also reached significance (*p* =.026). Intriguingly, the coefficient *β*_5_ which reflects the net sequential effect on latent decision tendency showed no effects in both conditions (*p*s >.555) and the between-condition comparison (*p* =.495). These net effects on latent decision are in line with standard behavioral observations of serial dependence (i.e., the point of subjective equality, see Appendix).

## Discussion

In the present study, we interleaved duration reproduction and bisection tasks to examine whether cross-task serial dependence in time perception stems from shared representations or is constrained by task-specific motor responses. Our results showed strong sequential biases within each task. In the reproduction task, participants exhibited a repulsion effect in perceptual serial dependence and an attraction effect in decisional carryover. Similarly, in the bisection task, we observed a repulsion in latent decision propensity and positive decisional carryover, consistent with previous findings (Li et al., 2023; Wiener et al., 2014). However, these serial dependencies disappeared when tasks alternated, indicating that shared duration encoding alone is insufficient to drive cross-task sequential biases. These findings highlight that response modes play a critical role in mediating task-specific serial dependence.

The perceptual repulsive bias observed in our within-task condition replicates findings from previous studies (Moon & Kwon, 2022; Sadil et al., 2024). Such repulsive bias is typically attributed to low-level sensory processing, such as neural adaptation (Moon & Kwon, 2022) or efficient encoding mechanisms that amplify differences between stimulus representations (Wei & Stocker, 2015). However, many studies also demonstrated attractive serial dependence (Cheng, Chen, & Shi, 2024; Cheng, Chen, Glasauer, et al., 2024; Cheng, Chen, Yang, et al., 2024; Glasauer & Shi, 2022; Wang et al., 2023), including the seminal study (Fischer & Whitney, 2014). The co-occurrence of repulsive and attractive effects complicates interpretation when using traditional one-factor models, which typically estimate net serial dependence - a mixture of opposing biases from different processing stages (Fritsche et al., 2020; Moon & Kwon, 2022).

By contrast, Structural Equation Modeling (SEM) enables us to separate sensory-level effects (direct repulsion from prior stimulus) from decision-level effects (attraction toward prior response), as portrayed in Figure 2. Specifically, our SEM framework models both direct pathways from prior stimulus to current report (capturing perceptual repulsion) and indirect pathways that mediate via prior response (capturing decision carryover). When summing these paths, we found that within-task net serial dependence becomes *attractive* in reproduction tasks (0.038 ± 0.013 in RR reproduction). This mirrors classic one-factor estimates but importantly hides the underlying sensory repulsion. Indeed, Sadil et al. (2024) cautioned that one-factor models can be misleading because they collapse distinct mechanisms into a single summary metric.

Even conventional regression analyses that include both prior stimulus and prior response often suffer from *multicollinearity* (as two factors are highly correlated), making it difficult to separately estimate repulsive and attractive contributions. In our work, we also conducted such regression analyses (see Appendix) and observed results largely aligned with but also diverged from SEM outcomes. For instance, conventional regression sometimes suggested ‘attractive’ perceptual serial dependence in reproduction – likely because it cannot model the indirect path via decision carryover, and the positive prior-response effect masks repulsion. The capacity of SEM to model latent constructs, measurement error, and causal pathways allows it to disentangle repulsive sensory effects that conventional methods miss.

The repulsive effect we observed within the bisection task also replicates earlier findings (Li et al., 2023). However, it’s crucial to note that this ‘repulsion’ occurs at the level of the latent decision propensity, not directly in the observed categorical responses. In other words, while prior durations shift the underlying probability of responding “long” away from the previous stimulus, this does not necessarily translate into final choice-level repulsion. Indeed, in our data the point of subjective equality (PSE), which indexes the actual bisection threshold, remained unchanged across prior task conditions (see Appendix). Therefore, caution is warranted: our probabilistic choice model reveals repulsion in latent decision tendencies, but this should not be equated with behavioral repulsion in overt judgments. The repulsive effect at the latent level may influence underlying discrimination sensitivity without affecting the observable bisection performance.

The key finding of the current study is the response-mode-specific serial dependence. At first glance, this may resemble previous findings that show no sequential effects across tasks (e.g., across numerosity and duration tasks in Fornaciai et al., 2023; Togoli et al., 2021) or across modalities (Li et al., 2023). However, duration representation likely differs from the representation of the other dimension (e.g., numerosity) or is encoded in a modality-specific manner at the low-level physiological processing. Since both task-related features (e.g., duration and numerosity) can be maintained simultaneously in working memory, previous reports of task-specific serial dependence may be attributed to an early-stage dimension- or modality-specific processing pathway (Togoli et al., 2021), occurring before the integration of sensory likelihood and prior knowledge in memory (Körding & Wolpert, 2004; Weiss et al., 2002). However, unlike these previous studies, the present study employed the same duration within the same visual modality for both timing tasks, ruling out this interpretation. Instead, our findings highlight response-specific serial dependence, suggesting that a separate, high-level processing stage occurs after early-stage duration encoding. The mechanisms underlying different consecutive task combinations (i.e., RR vs. JR and JJ vs. RJ) may probably differ in high-level processing after duration encoding, likely due to the reactivation of response-related tasks. Considering that both tasks exhibit attractive decisional carryover, it is more plausible that observers engage distinct response-dependent high-level processing for duration reproduction and bisection, making sequential effects from cross-task transitions less influential.

Previous research suggests that processing current inputs can reactivate memory of prior experiences (Bae & Luck, 2019; St. John-Saaltink et al., 2016), which may then integrate with ongoing memory representations (Zhang & Lewis-Peacock, 2024). Importantly, the strength of this memory reactivation has been linked to the strength of sequential bias (Barbosa & Compte, 2020). It is therefore possible that retrieval of task-related responses only reactivates prior duration with the same response mode in working memory. This is consistent with the theory of binding and retrieval in action control (BRAC, Frings et al., 2020), which suggests that stimulus features and responses are bound together as an event-file and retrieving the same response will activate the whole event file (Allenmark et al., 2025; Hommel, 2004). This could also explain previous findings of multiple motor-specific priors in interleaved duration reproduction and bisection tasks (Roach et al., 2017). Neuroimaging evidence further supports this idea. Studies have shown topographically organized duration maps in the human supplementary motor area (Kulashekhar et al., 2021; SMA, Protopapa et al., 2019), tuned to temporal contexts. The SMA is crucial for motor preparation and time perception (Coull et al., 2011; Kotz & Schwartze, 2011; Merchant et al., 2013). Additionally, sequential bias has been linked to activities in the left middle frontal gyrus (Cheng, Chen, Glasauer, et al., 2024), a region associated with the fronto-striatal pathway critical for working memory and time perception (Matell et al., 2005). These behavioral and neural findings imply the possibility of response-related memory reactivations in duration reproduction and bisection tasks.

It is worth noting that the current findings do not contradict previous interpretations of cross-task serial dependence. Rather, they enhance our understanding of the locus of interactions with serial dependence, indicating that responses engage with prior experiences at a later stage after encoding. If recent prior history occurs at early duration encoding, we would anticipate significant cross-task sequential effects, considering that the two tasks shared the same encoding. However, our results show the differential cross-task sequential effects, differing from studies that showed an early sequential effect in orientation perception (Cicchini et al., 2021). Cicchini et al. (2021) attributed the cross-context early-stage serial dependence to a hierarchical predictive coding framework, which posits that context-biased prior expectations are fed back to lower levels of sensory processing, where the expectation error would further transmit through the perceptual cascade (Friston, 2009; Rao & Ballard, 1999; Summerfield & de Lange, 2014). In contrast, our study emphasizes that recent history biases are closely linked to task-related responses. Specifically, activating a task-relevant response triggers reactivation of response-feature-bound event files in working memory, resulting in sequential biases. However, prior histories associated with different response modes remain inactive, reducing cross-task sequential biases. This response-binding was also evident in decisional carryover effects we observed (i.e., the decision itself as ‘long’ or ‘short’), which replicated many previous findings (Cheng, Chen, Yang, et al., 2024; Li et al., 2023; J. Wehrman et al., 2023; J. J. Wehrman et al., 2020; Wiener et al., 2014). Making a ‘long’ or ‘short’ response in the previous trial, both in reproduction and bisection, attracts the current response in that direction.

In summary, our findings reveal that serial dependence in time perception is tightly linked to task-specific motor reactivation, while decisional carryover remains unaffected by tasks. Despite a shared duration representation, sequential biases emerged only when motor responses were repeated, suggesting that response and duration are bound together and reactivation of past experiences depends on the retrieval of task-relevant responses. Our findings highlight the pivotal role of motor response in shaping cross-task serial dependence.

