## Appendix for "Response-driven Serial Dependence: When Response Mode Consistency Matters More than Shared Memory Encoding"

Standard analyses measuring serial dependence in duration reproduction tasks face confounding effects from prior duration and response hysteresis (or decisional carryover), similar to those observed in visual features like orientation (Sadil et al., 2024). In the time reproduction task, participants hold a key for a set period, causing the presented duration to correlate strongly with both the reproduced time and the reproduction bias. These correlations introduce multicollinearity, complicating efforts to separate the influences of prior duration from those of prior responses.

Here, we applied conventional multiple linear regression (one-factor analysis) and the linear mixed model (LMM, two-factor analysis), for a comparison with the structural equation modeling (SEM) we used in the manuscript.

**1. The reproduction task performance**

***One-factor analysis (multiple linear regression)***

The central tendency and serial dependence in the reproduction task were estimated by multiple linear regression with the current (*T_n_*) and prior predictor (*T_n-1_*): *Bias_n_* = *α* × *T_n_* + *𝛽* × *T_n-1_* + *ε.* The slope *α* indicates the central tendency index and the slope *𝛽* indicates the serial dependence index (SDI). The estimated SDI (0.021 ± 0.009) differed significantly from zero, *t*(23) = 2.224, *p* = .036, Cohen’s *d* = 0.454, *BF_10_* = 1.680, suggesting an effect of serial dependence. For different prior task types, the estimated SDI for RR trials (0.045 ± 0.011) was significantly positive (**Figure S1a**), *t*(23) = 4.108, *p* < .001, Cohen’s *d* = 0.839, *BF_10_* = 73.714, indicating an attractive sequential bias; while the mean SDI for JR trials (-0.004 ± 0.011) failed to reach significance, *t*(23) = -0.353, *p* = .727, Cohen’s *d* = 0.072, *BF_10_* = 0.227. The SDI differences between RR and JR trials was also significant (**Figure S1b**), *t*(23) = 3.902, *p* = .001, Cohen’s *d* = 0.891, *BF_10_* = 46.972.

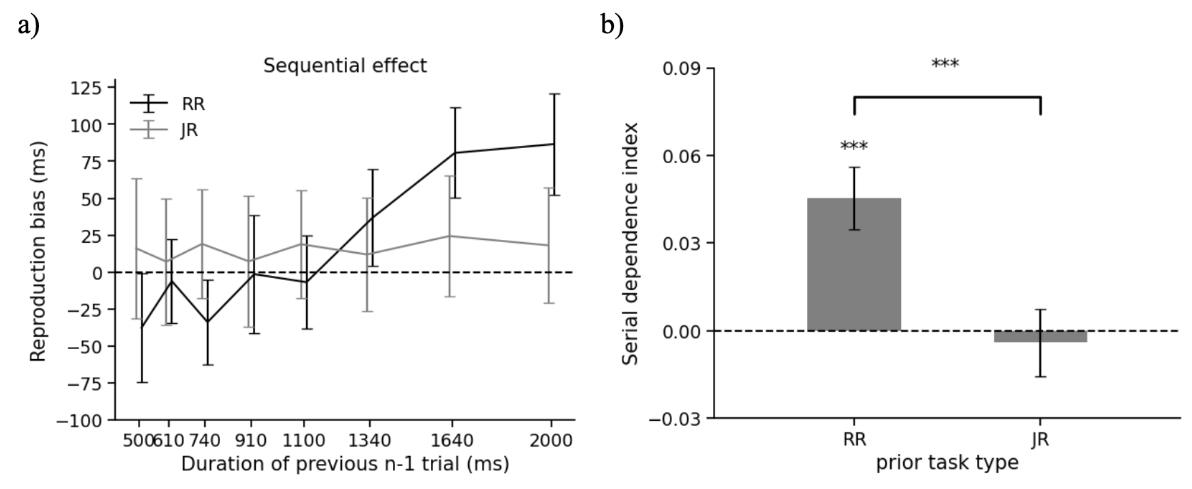

***Figure S1.*** *Sequential effect in the Reproduction task.* ***a)*** *the sequential effects based on previous duration in task-relevant RR and task-irrelevant JR trials. In particular, RR denotes the trials with a preceding task-relevant Reproduction task (Reproduction-Reproduction); while JR denotes the trials with a preceding task-irrelevant Judgment task (Judgment-Reproduction).* ***b)*** *the estimated serial dependence index based on previous duration in RR and JR trials. *** p* < .001*.*

***Two-factor analysis (linear mixed model)***

We examined fifteen multiple linear mixed-effects models (LMMs) for the reproduction task using the *lme4* package in R (see Table S1). In our model formulas, the dependent variable ***Bias_n_*** represents the current objective reproduction bias (i.e., reproduction - duration), while *T_n_* and *T_n-1_* mean the durations in the current and previous (*n*-1) trial, respectively. The factor ***Task_n-1_*** denotes the prior task type, either reproduction or judgment task. Additionally, we defined *Bias’_n_* as the subjective bias (i.e., reproduction - mean reproduction of the duration for each participant). ***J_n-1_*** denotes the prior judgment outcome (1=short, 2=long). ***Resp_n-1_*** reflects the subjective response of the prior duration being short or long; specifically, it is indexed by *J_n-1_* for the prior judgment task and by prior reproduction being shorter or longer than the mean reproduction of each participant (1=shorter, 2=longer) for the prior reproduction task.

As shown in Table S1, the best-fitting model (ranked by AIC) was model 14, which accounts for the current and prior durations, prior task type, prior reproduction bias and prior judgment, while also allowing random slopes and intercepts on the current and prior durations for each participant.

**Table S1 Comparisons between Linear Mixed-effect Models**

| **model** | **formula** | **AIC** | **BIC** |
| --- | --- | --- | --- |
| **model 14** | Bias_n_ ~ T_n-1_*Task_n-1_ + T_n_*Task_n-1_ + Bias_n-1_ + J_n-1_ + (T_n-1_+T_n_\|\|Sub) | 710.310 | 793.232 |
| **model 13** | Bias_n_ ~ T_n-1_*Task_n-1_ + T_n_*Task_n-1_ + Bias_n-1_ + (T_n-1_+T_n_\|\|Sub) | 717.966 | 793.978 |
| **model 15** | Bias_n_ ~ T_n-1_ + T_n_ + T_n-1_:Task_n-1_ + T_n_:Task_n-1_ + Bias_n-1_ + J_n-1_ + (T_n_+T_n-1_\|\|Sub) | 719.278 | 795.290 |
| **model 9** | Bias_n_ ~ T_n-1_*Task_n-1_ + T_n_*Task_n-1_ + Bias_n-1_ + (T_n_\|\|Sub) | 741.123 | 810.224 |
| **model 8** | Bias_n_ ~ T_n-1_*Task_n-1_ + T_n_*Task_n-1_ + Resp_n-1_ + (T_n_\|\|Sub) | 759.907 | 829.009 |
| **model 12** | Bias_n_ ~ T_n-1_ + T_n_ + T_n-1_:Task_n-1_ + T_n_:Task_n-1_ + Bias’_n-1_ + J_n-1_ + (T_n_+T_n-1_\|Sub) | 832.384 | 929.126 |
| **model 11** | Bias_n_ ~ T_n-1_*Task_n-1_ + T_n_*Task_n-1_ + Bias’_n-1_ + J_n-1_ + (T_n_+T_n-1_\|Sub) | 834.290 | 937.943 |
| **model 7** | Bias_n_ ~ T_n-1_*Task_n-1_ + T_n_*Task_n-1_ + (T_n_+T_n-1_\|Sub) | 837.338 | 927.170 |
| **model 4** | Bias_n_ ~ T_n-1_*Task_n-1_*T_n_ + (T_n_\|Sub) | 842.650 | 925.572 |
| **model 10** | Bias_n_ ~ T_n-1_*Task_n-1_ + T_n_*Task_n-1_ + Bias’_n-1_ + J_n-1_ + (T_n_\|\|Sub) | 863.466 | 939.478 |
| **model 5** | Bias_n_ ~ T_n-1_*Task_n-1_ + T_n_*Task_n-1_ + (T_n_\|\|Sub) | 865.778 | 927.969 |
| **model 6** | Bias_n_ ~ T_n-1_*Task_n-1_ + T_n_*Task_n-1_ + (T_n-1_\|\|Sub) | 2273.124 | 2335.316 |
| **model 3** | Bias_n_ ~ T_n-1_*Task_n-1_ + T_n_*Task_n-1_ + (1\|Sub) | 2278.774 | 2334.056 |
| **model 2** | Bias_n_ ~ T_n_ + T_n-1_ + (1\|Sub) | 2292.769 | 2327.320 |
| **model 1** | Bias_n_ ~ T_n-1_ + (1\|Sub) | 2301.048 | 2328.688 |

Figure S2 shows the mean reproduction biases based on the current duration (i.e., the central tendency effect; **Fig. S2a**) and the prior duration (i.e., the sequential effect; **Fig. S2b**), separated by prior task (Reproduction vs. Bisection). The best-fitting LMM model (model 14 in Table S1) includes reproduction bias as a function of the fixed effects of current duration, prior duration, prior report (prior reproduction bias and prior judgment choice), and prior task (with the base set as the relevant reproduction task), along with interactions between prior task and both prior and current durations. The model also incorporates random effects of current and prior durations for each participant (see estimates in **Table S2**).

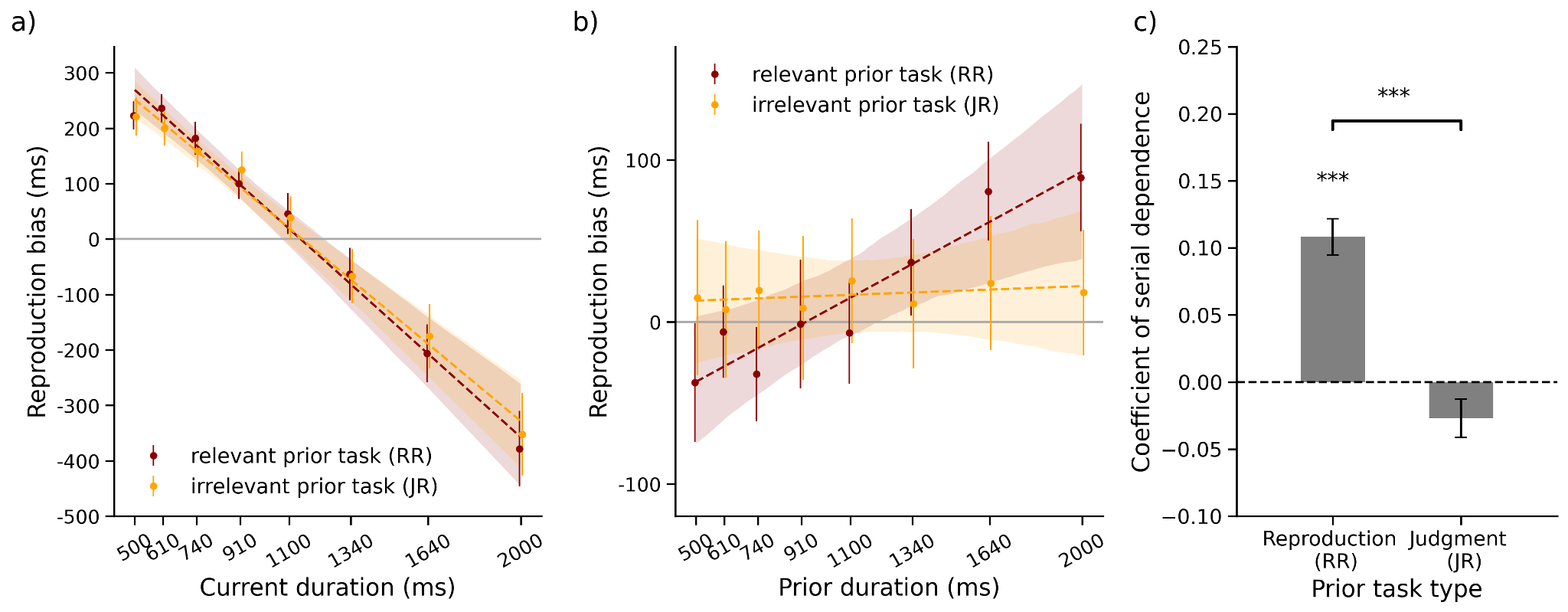

***Figure S2.*** *Reproduction task performance. (****a)*** *Reproduction bias as a function of current durations, shown for the prior task-relevant reproduction task (Reproduction-Reproduction; RR) and the prior irrelevant judgment task (Judgment-Reproduction; JR). (****b)*** *Reproduction bias as a function of previous durations, shown for prior task-relevant and -irrelevant conditions. (****c)*** *Mean coefficients of serial dependence for different prior task types, estimated by the best-fitting linear mixed effects (LMM) model. The error bar shows standard errors across participants. The dashed lines and color shades show linearly regressed predictions with 95% CI. *** p < .001.*

**Table S2.** Estimates for the fixed effects of the parameters were entered into the linear mixed-effects model to predict the current reproduction bias, with random effects of both the current and prior duration for each participant.

|  | **intercept** | **current duration** | **prior duration** | **prior bias**  **(R)** | **prior choice (J)** | **prior task**  **(J)** | **interaction** | |
| --- | --- | --- | --- | --- | --- | --- | --- | --- |
|  |  |  |  |  |  |  | **current dur. × prior task** | **prior dur. × prior task** |
| 𝛽 | 0.352 | -0.414 | 0.108 | 0.162 | 0.034 | 0.074 | 0.027 | -0.135 |
| ***t*** | 9.198 | -8.223 | 8.010 | 12.121 | 3.113 | 3.314 | 2.258 | -9.165 |
| ***p*** | < .001^***^ | < .001^***^ | < .001^***^ | < .001^***^ | .002^**^ | < .001^***^ | .024^*^ | < .001^***^ |

Note: The prior task was binary coded: prior Reproduction (R) = 0; prior bisection Judgment (J) = 1. Thus, the main coefficients were estimated for the prior reproduction condition, while the interactions were additive coefficients for correspondent interactive terms in the prior Judgment condition. *, **, and *** represent *p* < .05, .01, and .001 respectively.

To obtain statistical tests, we referenced the prior task to the Bisection Judgment and conducted another LMM estimation using the best model. Table S3 presents the coefficient results, which are essentially the same as those in Table S2. In this table, we obtained the slope of the prior duration for the previous trial for the judgment task condition.

**Table S3.** Estimates for the fixed effects of the parameters were entered into the linear mixed-effects model to predict the current reproduction bias, with random effects of both the current and prior duration for each participant.

|  | **intercept** | **current duration** | **prior duration** | **prior bias**  **(R)** | **prior choice (J)** | **prior task**  **(R)** | **interaction** | |
| --- | --- | --- | --- | --- | --- | --- | --- | --- |
|  |  |  |  |  |  |  | **current dur. × prior task** | **prior dur. × prior task** |
| 𝛽 | 0.426 | -0.387 | -0.027 | 0.162 | 0.034 | -0.074 | -0.027 | 0.135 |
| ***t*** | 9.878 | -7.697 | -1.902 | 12.121 | 3.113 | -3.314 | -2.258 | 9.165 |
| ***p*** | < .001^***^ | < .001^***^ | .061^†^ | < .001^***^ | .002^**^ | < .001^***^ | .024^*^ | < .001^***^ |

Note: The prior task was binary coded: prior Reproduction (R) = 1; prior bisection Judgment (J) = 0. Thus, the main coefficients were estimated for the prior judgment condition, while the interactions were additive coefficients for correspondent interactive terms in the prior reproduction condition. *, **, and *** represent *p* < .05, .01, and .001 respectively, and ^†^ marginal significance.

The serial dependence was significantly positive for the RR condition (𝛽 = 0.108, *p* < .001, Table S2) but only marginally negative for the JR condition (𝛽 = -0.027, *p* = .061, Table S3). Permutation tests on sequential effects, however, revealed both coefficients fell beyond the 95% CIs of the null distribution (1000 shuffling iterations, RR: [-0.020, 0.018]; JR: [-0.024, 0.022]), though the latter was close to the lower boundary, consisting with the LMM results. Prior durations biased current duration reproduction in the same direction for 10.8% if both responses were reproduction. The prior judgment task induced a mild repulsion effect. The decision carryover effects, defined by prior bias or prior choice, were significant for both (prior reproduction bias: 𝛽 = 0.162, *p* < .001; prior choice: 𝛽 = 0.034, *p* = .002 with Short coded as 1, Long as 2). Additionally, the prior judgment task increased the current reproduction by 74 ms in general (*p* < .001). Importantly, the prior judgment task affected the central tendency and serial dependence differently than the prior reproduction task, as shown by the interaction between the prior task and the prior and current durations (*p*s < .024).

Taken together, the one-factor analysis observed an attractive sequential effect from prior duration in task-relevant prior trial type (RR), but mixed with the effect from prior reproduction. The two-factor analysis observed attractive effects from both predictors. Though segregated the effects from prior duration and prior reproduction, it introduced the problem of multicollinearity, that is, the correlation between the predictors (prior duration and prior response).

**2. The judgment task performance**

***Behavioral observation of serial dependence based on PSE***

In the judgment task, we evaluated the sequential effect based on previously-perceived duration. We estimated the *point of subjective equality* (*PSE*) for preceding short and long durations (i.e., duration for the previous *n*-1 trial shorter or longer than the standard 1000 ms). Sequential effects would have occurred if the PSEs between trials preceded by short and long durations differed from each other. As a result, the overall sequential effect was not significant, *t*(23) = -0.382, *p* = .706, Cohen’s *d* = 0.045, *BF_10_* = 0.229, with comparable mean PSEs for trials with previous short (1039 ± 39) and long (1048 ± 44) duration. Moreover, the sequential effect was non-significant for both transitional task relevances [JJ: *t*(23) = 0.387, *p* = .702, Cohen’s *d* = 0.052, *BF_10_* = 0.230; RJ: *t*(23) = -0.543, *p* = .592, Cohen’s *d* = 0.121, *BF_10_* = 0.245]. These results suggest that the prior duration had little bearing on the current judgment, regardless of its task relevance.

***Serial dependence by the*** ***probabilistic choice model***

We further estimated the sequential effect using the *probabilistic choice model* (Feigin, Baror, et al., 2021; Feigin, Shalom-Sperber, et al., 2021; Li et al., 2023). Specifically, the probability of making a ‘long’ response is assumed to be predicted by the binomial logistic regression with a decision variable z: $p_{t}(long)=\frac{1}{1+e^{-z_{n}}}$ , where the decision variable (*z_n_*) is influenced by the current (*T_n_*), the prior duration (*T_n-1_*), the prior report (PR_n-1_: Short vs. Long), the prior task (*Task_n-1_*: Reproduction vs. Judgment), the prior judgment (*J_n-1_*), and their possible interactions, similar to the LMMs used for the reproduction task. A slight variation was that previous biases in the reproduction task were transformed into binary encoded labels of ‘Short’ and ‘Long’, aligning with the probabilistic choice model. Additionally, we considered both the original linear-scaled (*T*) and log-scaled (i.e., log(*T*)) current and previous durations in the LMMs. It turned out that the log-scaled LMMs fitted better for the judgment task.

***Serial dependence based on LMM***

We analyzed performance on the binary judgment task performance using multiple linear mixed-effects models (LMMs) with logistic regression (i.e., binomial family) in the *lme4* package in R. In our model formulas, the dependent variable ***J_n_*** in the formula denotes the current binary judgment (0=short, 1=long). ***T_n_*** and ***T_n-1_*** indicate the durations of the current and previous (*n*-1) trials, respectively. Following the literature (Feigin, Baror, et al., 2021; Feigin, Shalom-Sperber, et al., 2021; Li et al., 2023), we also transformed the durations into the log-scale and normalized with the root mean square (RMS), denoted by ***LT_n_*** and ***LT_n-1_*** for the current and previous duration respectively. We considered both the original linear-scaled and log-scaled models. By testing multiple LMMs, differing only in the scales, we found that the log-scaled models were generally better than the linear-scaled models. Thus, in the listed compared models (Table S4), we primarily included the log-scaled models (linear-scaled models were Model 8 and 9).

The factor ***Task_n-1_*** denotes the prior task type, either judgment or reproduction task. ***Resp_n-1_*** depicts the subjective response of the prior duration being short or long. Specifically, for the prior judgment task, it corresponds to a ‘short’ vs. ‘long’ response, while for the prior reproduction task, it is determined by whether the prior reproduction was shorter or longer than the mean reproduction of each participant. The prior responses were recorded as -1 (short) and 1 (long). In some models, we used prior reproduction bias (*Bias_n-1_*: reproduction - mean reproduction of the duration for each participant) and prior judgment (*J_n-1_*: 1=short, 2=long), separately. As shown in Table S4, the best-fitting model (ranked by AIC) was Model 13 with parameters of the current and prior durations, prior task type, prior response, while allowing random slopes and intercepts on the current and prior durations for each participant.

**Table S4**

| **model** | **formula** | **AIC** | **BIC** |
| --- | --- | --- | --- |
| **model 13** | J_n_ ~ LT_n_*Task_n-1_ + LT_n-1_*Task_n-1_ + Resp_n-1_*Task_n-1_ + (LT_n_ + LT_n-1_\|Sub) | 5725.502 | 5822.165 |
| **model 14** | J_n_ ~ LT_n_*Task_n-1_ + LT_n-1_*Task_n-1_ + Resp_n-1_ + (LT_n_ + LT_n-1_\|Sub) | 5730.554 | 5820.312 |
| **model 12** | J_n_ ~ LT_n_*Resp_n-1_ + LT_n-1_*Resp_n-1_ + (LT_n_ + LT_n-1_\|Sub) | 5731.026 | 5813.880 |
| **model 16** | J_n_ ~ LT_n_*Task_n-1_ + LT_n-1_*Task_n-1_ + J_n-1_ + Bias_n-1_ + (LT_n_ + LT_n-1_\|Sub) | 5731.222 | 5827.885 |
| **model 7** | J_n_ ~ LT_n_*Task_n-1_ + LT_n-1_*Task_n-1_ + (LT_n_ + LT_n-1_\|Sub) | 5758.973 | 5841.827 |
| **model 8** | J_n_ ~ T’_n_*Task_n-1_ + T’_n-1_*Task_n-1_ + (T’_n_\|\|Sub) | 5867.928 | 5923.164 |
| **model 15** | J_n_ ~ LT_n_*Task_n-1_ + LT_n-1_*Task_n-1_ + Resp_n-1_*Task_n-1_ + (LT_n_\|\|Sub) | 5870.126 | 5939.171 |
| **model 11** | J_n_ ~ LT_n_ + LT_n-1_ + Resp_n-1_ + (LT_n_\|\|Sub) | 5873.664 | 5915.091 |
| **model 6** | J_n_ ~ LT_n_*Task_n-1_ LT_n-1_*Task_n-1_ + + (LT_n_\|\|Sub) | 5900.675 | 5955.911 |
| **model 5** | J_n_ ~ LT_n_*LT_n-1_*Task_n-1_ + (LT_n_\|\|Sub) | 5903.205 | 5972.250 |
| **model 9** | J_n_ ~ T’_n_*Task_n-1_ + T’_n-1_*Task_n-1_ + (T’_n_ + T’_n-1_\|Sub) | 5990.488 | 6073.342 |
| **model 10** | J_n_ ~ LT_n_ + LT_n-1_ + Resp_n-1_ + (1\|Sub) | 6094.135 | 6128.657 |
| **model 1** | J_n_ ~ LT_n_ + (1\|Sub) | 6116.216 | 6136.929 |
| **model 2** | J_n_ ~ LT_n_ + LT_n-1_ + (1\|Sub) | 6118.205 | 6145.823 |
| **model 3** | J_n_ ~ LT_n_*LT_n-1_ + (1\|Sub) | 6119.715 | 6154.238 |
| **model 4** | J_n_ ~ LT_n_*LT_n-1_*Task_n-1_ + (1\|Sub) | 6124.831 | 6186.971 |

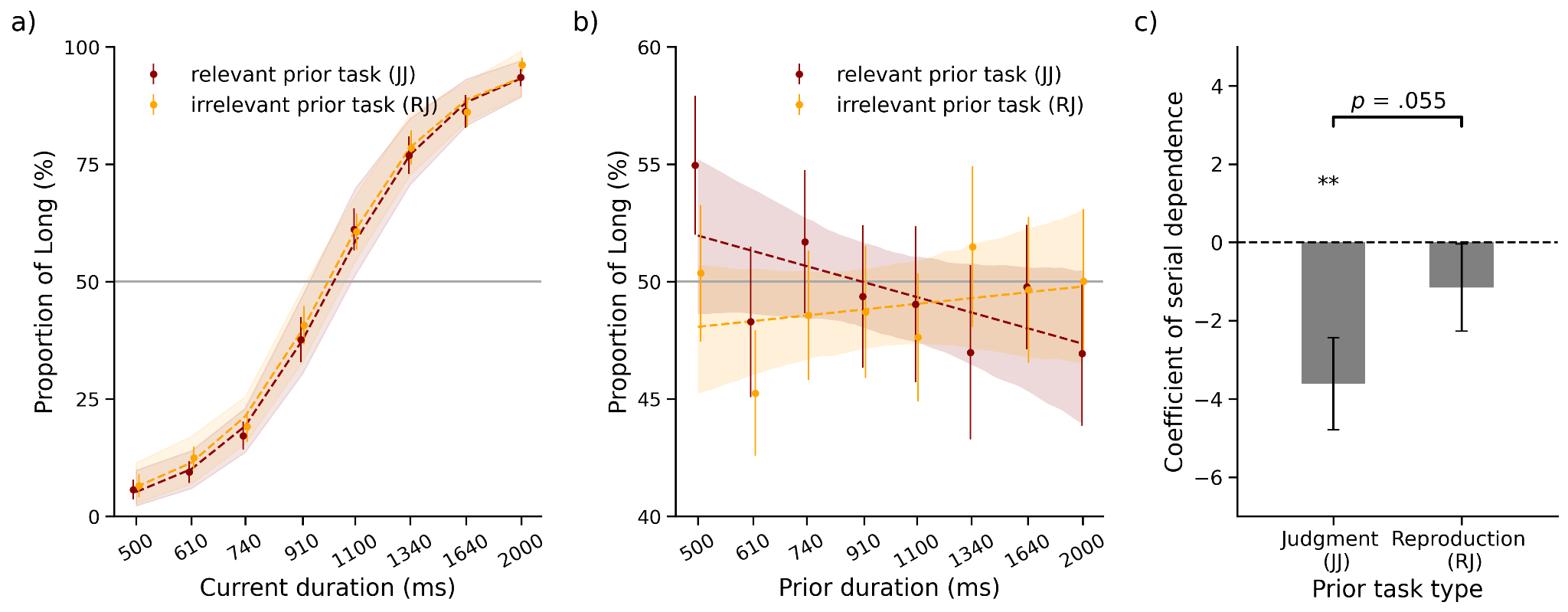

***Figure S3.*** *Judgment task performance. (****a)*** *The proportion of long reports (PL) as a function of current durations with prior task-relevant reproduction tasks (Judgment-Judgment or JJ) and prior irrelevant judgment tasks (Reproduction-Judgment or RJ). The dashed curves and color shades show the predictions from the selected best model with 95% CI. (****b)*** *The PLs as a function of previous durations with prior task-relevant and -irrelevant tasks. The dashed lines and shades show simple linear fitting with 95% CI based on the log-scale of prior durations. (****c)*** *Prior task effect of perceptual serial dependence: the estimated coefficients of prior duration in different prior task types by the best-fitting model. The error bar shows the standard error of participants. ** p < .01.*

**Table S5.** Estimations for the fixed effects of the parameters entered in the logistic regression model predicting the current judgment choice, with random effects of both the current and prior duration for each participant.

|  | **intercept** | **current duration**  **(log)** | **prior duration**  **(log)** | **prior report** | **prior task**  **(R)** | **interaction** | | |
| --- | --- | --- | --- | --- | --- | --- | --- | --- |
|  |  |  |  |  |  | **current dur. (log) × prior task** | **prior dur. (log) × prior task** | **prior report × prior task** |
| 𝛽 | -34.888 | 38.486 | -3.612 | 0.368 | -0.572 | -1.770 | 2.465 | -0.228 |
| ***Z*** | -10.812 | 12.391 | -3.059 | 5.651 | -0.322 | -1.326 | 1.922 | -2.664 |
| ***p*** | < .001^***^ | < .001^***^ | .002^**^ | < .001^***^ | .748 | .185 | .055^†^ | .008^**^ |

Note: The current and prior duration were transformed in the log-scaled and normalized with the related root mean square (RMS), following a similar approach used by Li and colleagues (2023). The prior tasks were binary-coded: Prior Judgment (J) = 0 and Prior Reproduction (R) = 1. The main coefficients estimate effects for the Prior Judgment condition, while interaction terms represent additive effects in the Prior Reproduction condition for correspondent interactive terms. Prior reports were recoded as -1 for “Short” and 1 for “Long”. **, ***, and † represent *p* < .01, <.001, and marginal significance, respectively.

Analogy to previous analysis, we referenced the prior task to the Reproduction task and conducted the best-fitting LMM model. Table S6 shows the counterpart of Table S5. The slope of the prior duration for the reproduction task was shown in the table.

**Table S6.** Estimations for the fixed effects of the parameters entered in the logistic regression model predicting the current judgment choice, with random effects of both the current and prior duration for each participant.

|  | **intercept** | **current dur.**  **(log)** | **prior dur.**  **(log)** | **prior report** | **prior task**  **(J)** | **interaction** | | |
| --- | --- | --- | --- | --- | --- | --- | --- | --- |
|  |  |  |  |  |  | **current dur. (log) × prior task** | **prior dur. (log) × prior task** | **prior report × prior task** |
| 𝛽 | -35.404 | 36.668 | -1.154 | 0.141 | -0.572 | 1.778 | -2.421 | 0.226 |
| ***Z*** | -11.005 | 11.882 | -1.033 | 2.447 | -0.322 | 1.332 | -1.892 | 2.646 |
| ***p*** | < .001^***^ | < .001^***^ | .301 | .014^*^ | .748 | .183 | .059^†^ | .008^**^ |

Note: the prior task was binary coded: prior Judgment (J) = 1; prior Reproduction (R) = 0. Thus, the main coefficients were estimated for the prior reproduction condition, while the interactions were additive coefficients for corresponding interactive terms in the prior judgment condition. Prior report was recoded as -1 if it was “Short” and 1 if “Long”. ** and *** represent *p* < .01, and .001, respectively, and ^†^ marginal significance.

The results showed the current duration dominated the bisection decision (𝛽 = 38.486, *p* < .001, **Fig. S3a**), with no significant difference between the two prior tasks (*p* = .185). The prior duration negatively impacted the current bisection task (i.e., repulsion, 𝛽 = -3.612, *p* = .002) when the prior task was the same (i.e., JJ), and the impact was reduced to non-significance (𝛽 = -1.154 ± 1.181, *p* = .301) in the cross-task RJ condition. The reduction was marginally significant (*p* = .055, **Fig. S3b**). The results were further confirmed by permutation tests (1000 shuffling iterations), which showed the coefficient in the JJ condition fell beyond the 95% CIs of the ‘null’ distribution ([-1.777, 1.866]), but not the RJ condition ([-1.603, 1.493]), similar to the findings of differential cross-task effects shown in the reproduction task.

The decision carryover was also significant for both the prior judgment task (𝛽 = 0.368, *p* < .001, Table S5) and the prior reproduction task (𝛽 = 0.141, *p* = .014, Table S6), with a significant difference between two prior tasks (*p* = .008). Like the reproduction task, prior decisions had a positive impact on the current judgments.

To summarize, the probabilistic choice model estimated a repulsive serial dependence from prior duration but an attractive serial dependence from prior response.
